# Study of the Effects of Different Spray Adjuvants on Unmanned Aerial Vehicle Crop Protection in Rice Fields

**DOI:** 10.64898/2025.12.09.693308

**Authors:** Weicai Qin, Xiaotuo Wang, Xiaolan Lv

## Abstract

Unmanned aerial vehicle (UAV) spraying is being increasingly used for rice pest control, but challenges remain in terms of optimizing the uniformity and efficacy of droplet deposition. In this study, the effects of six commercial spray adjuvants on the UAV-based control of rice stem borers in paddy fields were systematically evaluated. With a T30 UAV equipped with water-sensitive paper monitoring, we assessed the droplet spreading dynamics, deposition density, coefficient of variation (CV), and field control efficacy at various heights (1.5–3 m) and speeds (3–6 m/s). The results revealed that Activator 90 significantly reduced the CV by 30% (p<0.01) and resulted in the highest deposition density (30.0 drops/cm^2^) within 2 minutes. Compared with the control treatment, the Activator 90 increased the control efficacy by 70% to 87%, with optimal parameters at a height of 2.5 m and a spraying rate of 22.5 L/ha. This research reveals that selective adjuvants can markedly improve UAV spray performance, providing a practical framework for precision pest management in rice ecosystems.

## 1. Introduction

The challenges of drift, evaporation, and poor leaf adhesion of pesticide droplets during spraying significantly reduce the effective coverage and pesticidal efficacy, limiting the utilization rate of applied chemicals^1-3^. In 2015, the Ministry of Agriculture and Rural Affairs launched a zero-growth plan for the use of fertilizers and pesticides, with the goal of controlling the amount of fertilizers and pesticides used within a specific range to promote sustainable development in agriculture^4-6^. In 2016, the total amount of pesticides used in China was 1.7831 million tons per year, a decrease of 5.56% compared with the previous year. In January 2021, the Ministry of Agriculture and Rural Affairs issued another document, stating that it will strive to achieve a pesticide utilization rate of 90% by 2025, with zero growth in the use of chemical pesticides^7-9^.

Owing to the low efficiency of manual pesticide spraying and the substantial environmental pollution caused by excessive pesticide use, the combination of agricultural unmanned aerial vehicles and spray adjuvants has achieved precise, efficient, and low-cost pesticide application^10-14^. He et al. investigated the effects of spray adjuvants and application volume on the deposition density, deposition amount, and effective deposition rate of droplets at different positions in the rice canopy. The results of an unmanned aerial vehicle spraying experiment revealed that the addition of spray adjuvants can significantly increase the deposition amount and effective deposition rate of droplets^15^. Lan et al. investigated the effects of spray adjuvants on the surface tension of the solution, the contact angle of the medicine on corn leaves, and the deposition characteristics of droplets and compared the deposition characteristics of different adjuvants sprayed by unmanned aerial vehicles^16^. Han et al. investigated the distribution of deposition amounts on the canopies and ground of hilly citrus trees under the precision operation mode of agricultural unmanned aerial vehicles and the effect of different spray adjuvants^17^. Guo et al. conducted pesticide spraying experiments using unmanned aerial vehicles to analyse the effects of different adjuvants on the distribution of droplet deposition on the rice canopy and the effect of adjuvant-assisted unmanned aerial vehicles in controlling rice blast disease^18^. Wang et al. compared the performance of various types of tank-mixed adjuvants by analysing the droplet profiles, drift potential index (DIX), field deposition and control of wheat rust and aphids in a wind tunnel to elucidate the spray characteristics and control of these adjuvants in aerial sprays^19^. Yan et al. explored the effects of aerial spray adjuvants on droplet deposition and distribution in wheat and tested the research hypothesis that the use of aerial spray adjuvants could reduce wheat mycotoxin contamination while improving Fusarium head blight of wheat control^20^.

However, most previous studies have focused on either individual adjuvants or specific crop types, with limited comparative research on standardized UAV platforms using multiple spray adjuvants under uniform field conditions. In particular, systematic evaluations of the effects of commonly used adjuvants on droplet deposition characteristics, spray coverage, and pest control efficacy in rice paddies are scarce. While these adjuvants are extensively applied in agricultural practices, their relative performance under UAV-based spraying conditions, especially for controlling the rice stem borer, remains ambiguous. Therefore, this study aims to fill this research gap by systematically evaluating six commercial adjuvants through integrated laboratory and field trials, with the specific objectives of (1) characterizing their dynamic spreading behaviour on rice leaves; (2) quantifying their effects on UAV spray droplet deposition density and uniformity; (3) determining optimal operational parameters for effective spray width when the best-performing adjuvant is used; and (4) evaluating field control efficacy against a rice stem borer. We hypothesized that compared with nonadjuvant treatments, adjuvant treatments would significantly affect droplet spreading characteristics, improve deposition quality while improving uniformity, increase effective spray width under optimized parameters, and ultimately provide superior pest control efficacy. The findings are expected to establish selection criteria and operational guidelines for precision UAV pesticide application in rice production systems.

## 2. Materials and methods

### 2.1. Materials

The experiments were conducted in the National Modern Agricultural Demonstration Zone of Wujiang, Jiangsu Province, which is very suitable for the growth of crops such as wheat, rice, rapeseed, and aquatic plants. The terrain is flat without mountains, making full mechanized management easy throughout the process^21^. The rice variety used was Nanjing 46, which was transplanted by machine and had a plant height of 25–35 cm. The tested pesticides and varieties were Kangkuan (20% chlorfenapyr suspension concentrate) at a dose of 150 mL/ha and the spray-specific adjuvants Silwet L-77 (surfactant), Sylgard 309 (oil-based spreader-sticker), Activator 90 (surfactant/sticker), TM07 (deposition aid), LI-700 (spreading agent), and Aeromate 480 (drift reducer/adhesion promoter) at a dose of 150 mL/ha. These adjuvants were chosen on the basis of their suggest roles in modulating droplet behaviour, increasing foliar retention, and reducing drift under field conditions. This study was provided by the Institute of Plant Protection, Chinese Academy of Agricultural Sciences. The physical properties of the spray adjuvants are listed in Table 1.

**Table 1.** Physical property parameters.

| Adjuvant Name | Surface Tension (mN/m) | Viscosity (mPa·s) | Density (g/cm<sup>3</sup>) |
| --- | --- | --- | --- |
| Activator 90 | 20.5 | 1.05 | 0.98 |
| TM07 | 23.8 | 1.30 | 1.01 |
| Sylgard 309 | 22.0 | 1.10 | 1.00 |
| Aeromate 480 | 28.7 | 2.50 | 1.05 |
| Silwet L-77 | 18.9 | 0.95 | 1.00 |
| LI-700 | 30.5 | 2.80 | 1.06 |

This spraying operation uses a T30 plant protection unmanned aerial vehicle (UAV) manufactured by DJI Innovation Technology Co., Ltd. of Shenzhen. The specific parameters of the plant protection UAV are shown in Table 2. The T30 UAV body is constructed of lightweight materials and has high strength and rigidity. The UAV controller is the core control system of T30 and can adjust and monitor the parameters of plant protection operations. The UAV controller is equipped with multiple motors and propellers and provides powerful lifting and flight capabilities. GPS navigation technology is used to accurately locate and control the position and flight trajectory of the UAV. The dispensing system uses intelligent dispensing technology, which can automatically adjust the proportion of liquid medicine according to the needs of different crops. The structure of the DJI T30 is shown in Figure 1.

**Table 2.** Main parameters of the T30 plant protection UAV.

| Main parameters | Value |
| --- | --- |
| Dimensions/mm×mm×mm | 1180×1180×410 |
| Tank capacity/L | 6, 8, and 10 |
| Flight speed/m·s<sup>-1</sup> | 1–8 |
| Flight height/m | 1–10 |
| Spray width/m | 1.5-3 |
| Nozzle diameter/mm | 60 |
| Continuous operating hours/min | 20 |

**Figure 1.**
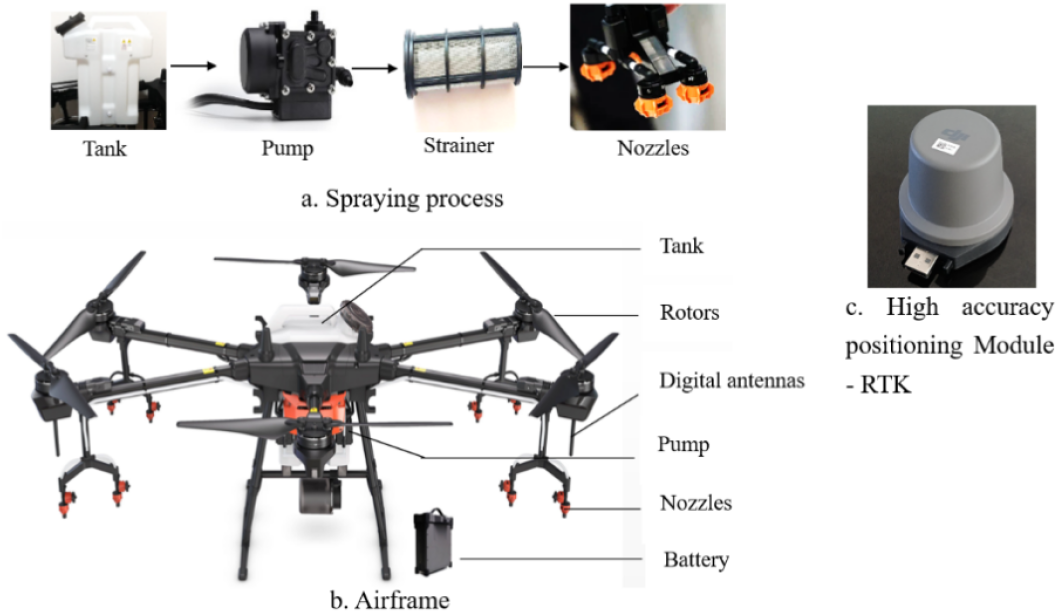
T30 plant protection UAV.

### 2.2 Methods

#### 2.2.1. Preliminary Evaluation of Spray Adjuvants

A 10 μL pipette was used to aspirate the solutions of Silwet L-77, Sylgard 309, Activator 90, TM07, LI-700, and Aeromate 480 separately. The liquid droplets were dropped onto the rice leaves, and the droplet spreading effect and spreading time were observed.

#### 2.2.2 Spray Droplet Deposition Test

In accordance with the actual field conditions, common parameters for the plant protection UAV were selected for testing the uniformity of mist droplet deposition and field efficacy in a 1.3 ha rice field, which was divided into 8 plots. The design parameters are listed in Table 3. Sampling was performed at five points in the upper part of the crop in each plot. The experimental machinery operation and sampling point layout are shown in Figure 2.

**Table 3.** Experimental design.

| Treatment code | Machine type | Operating height | Operating speed | Spray Volume | Pesticides and adjuvants | Area (ha) | Remarks |
| --- | --- | --- | --- | --- | --- | --- | --- |
| 1 | T30 Plant Protection UAV | 2.0 m | 4 m/s | 15 L/ha | Pesticides +Silwet L-77 | 0.20 | In each plot, five water-sensitive papers were randomly placed for field observation |
| 2 |  |  |  |  | Pesticides +Sylgard 309 | 0.21 |  |
| 3 |  |  |  |  | Pesticides +Activator 90 | 0.11 |  |
| 4 |  |  |  |  | Pesticides +TM07 | 0.29 |  |
| 5 |  |  |  |  | Pesticides +LI-700 | 0.25 |  |
| 6 |  |  |  |  | Clearwater | 0.09 |  |
| 7 |  |  |  |  | Pesticides | 0.09 |  |
| 8 |  |  |  |  | Pesticides +Aeromate 480 | 0.15 |  |

**Figure 2.**
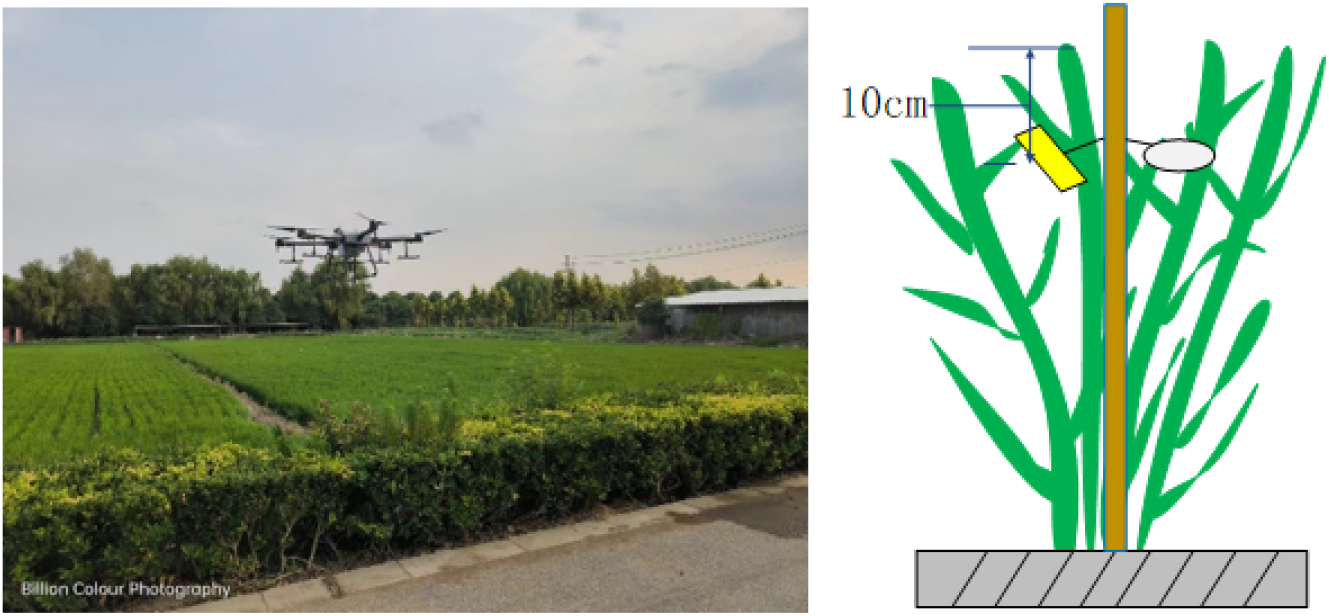
Machine operation and sampling point layout.

#### 2.2.3. Effective spray width test

The effects of four factors, including the flight height A (1.5 m, 2 m, 2.5 m, and 3 m), flight speed B (3 m/s, 4 m/s, 5 m/s, and 6 m/s), application rate per ha C (15 L/ha and 22.5 L/ha), and adjuvant Activator 90 (with an adjuvant and without an adjuvant), on the effective spraying width of the T30 plant protection UAV were investigated. The experimental process is shown in Figure 3, with a total of 15 poles arranged at intervals of 0.5 m, each with water-sensitive paper attached. The T30 plant protection UAV flew along the centerline, and the operation parameters are listed in Table 4.

**Table 4.**
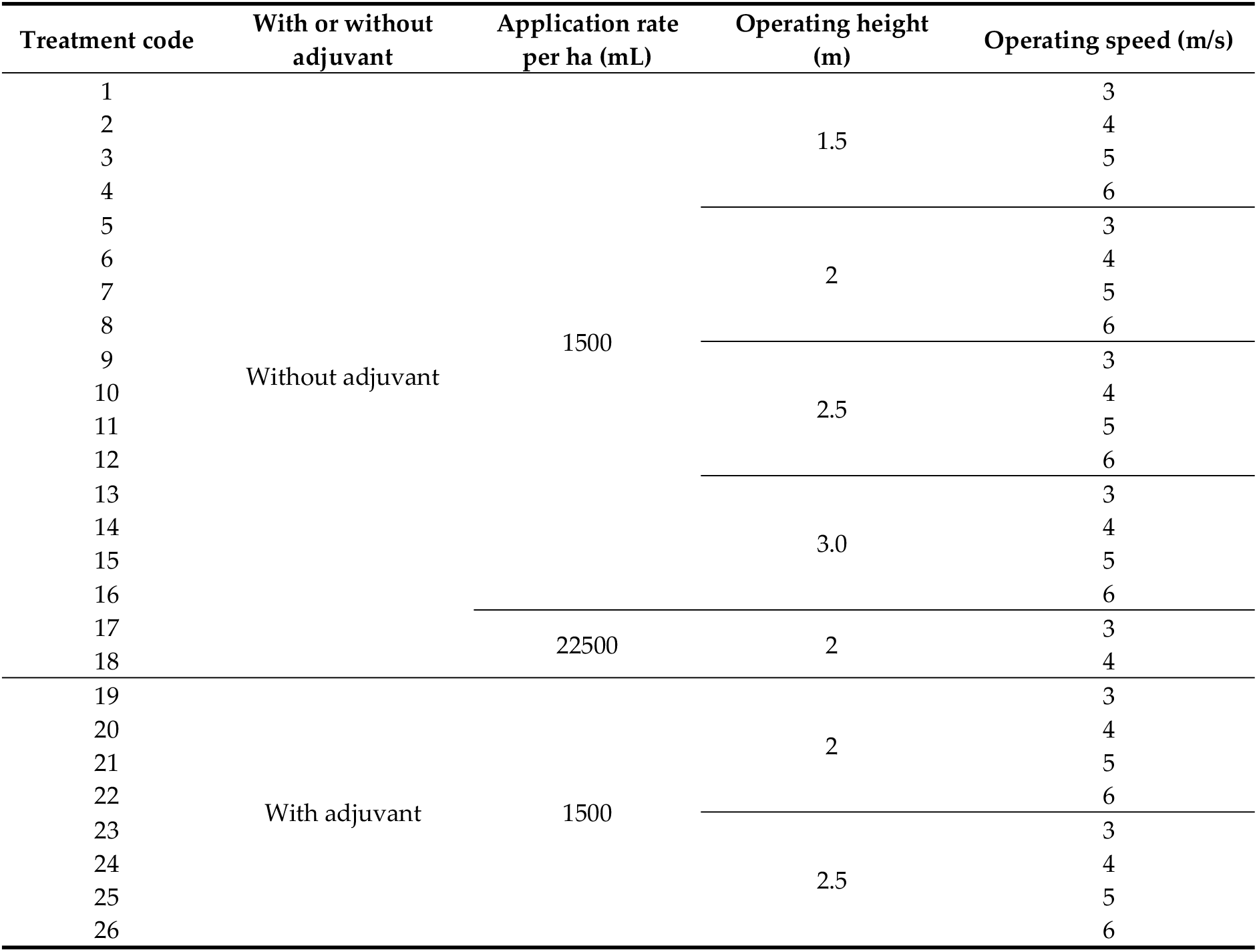
Design of effective spray width operating parameters.

**Figure 3.**
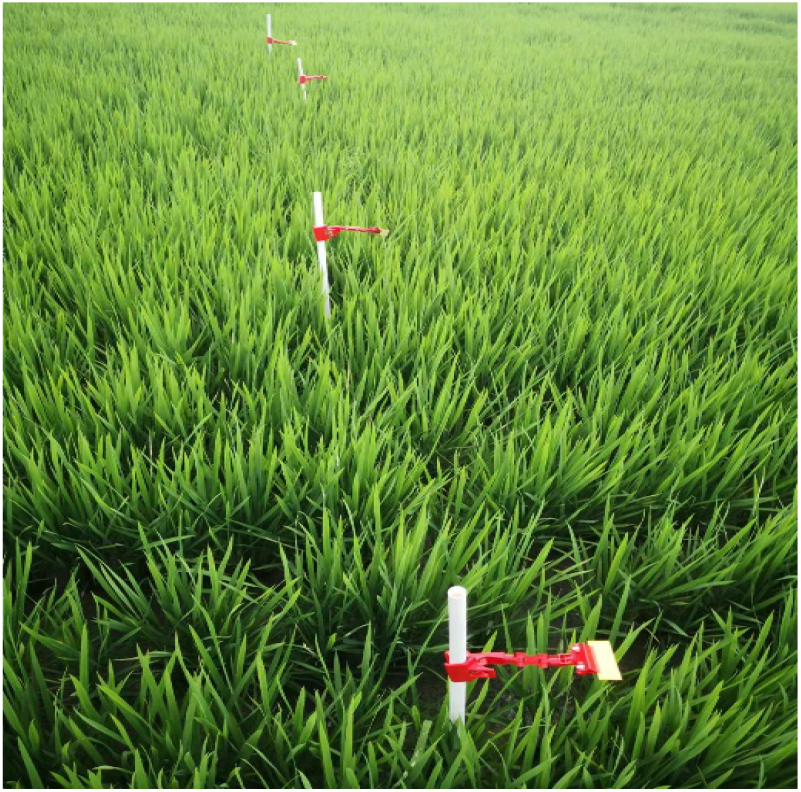
Layout of the effective spray width position.

#### 2.2.4. Pest control trials in the field

In accordance with the “Guidelines for Field Efficacy Trials of Pesticides Part 1: Insecticides for the Control of Rice Stem Borers (GB/T 17980.1—2000), the efficacy investigation was conducted on August 7 (7 days after spraying, tillering stage), August 14 (14 days after spraying, preflowering stage), and August 22 (21 days after spraying, mid-flowering stage)^22-23^. Ten sampling points were randomly selected in each plot, and each point was investigated for an area of 0.25 m^2^. The investigation encompasses 2–5 m^2^ of rice fields, and the number of tillers, blight plants, leaves, and affected plants (rolled leaves) were recorded^24-26^.

### 2.3. Data processing

After the experiment, the water-sensitive papers were collected, labelled, and sealed for preservation. The results are shown in Figure 4 and were returned to the laboratory for analysis. The water-sensitive papers were scanned using a scanner (resolution 600 dpi) and processed using the image processing software Photoshop for edge processing. A specific area was cropped and saved in JPG format and then analysed using DepositScan software^27-28^.

**Figure 4.**
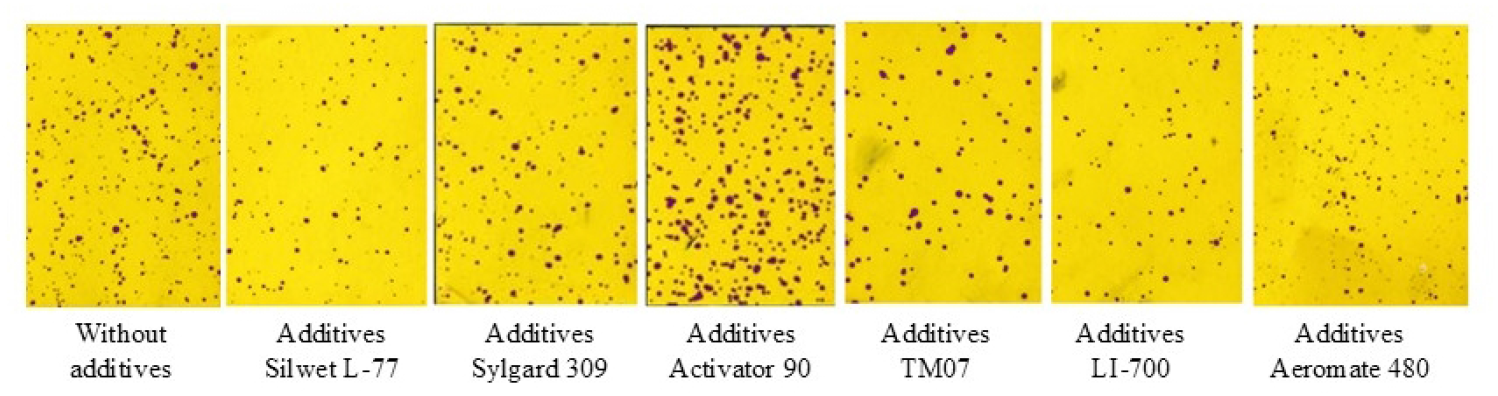
Distribution of droplets with different adjuvants.

The coverage rate on the sampling card can be obtained by calculating the ratio of the number of fog droplet pixels in the analysis area of the image to the analysis area. The calculation formula is^29-31^:

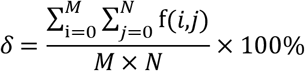

where M represents the width of the analysis area in pixels, N denotes the height of the analysis area in pixels, and f(i, j) indicates the grey value flag of the pixel in the image analysis area at coordinates (i, j) relative to the pixel. If the pixel is black, then f(i, j)=1; otherwise, f(i, j)=0.

The coefficient of variation V is used as a measure of the uniformity of the droplet distribution, where V is calculated using the following formula^32-33^:

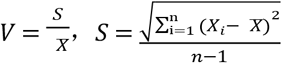

where S represents the standard deviation, X_i_ denotes the number of droplets per unit area of each sampling card, 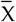 indicates the average number of droplets per unit area of the sampling card, and n represents the total number of sampling cards per layer.

In accordance with the industry standard “Technical Specification for Quality Evaluation of Agricultural Unmanned Aerial Vehicles”, a sedimentation density of ≥15 drops/cm^2^ is used as the boundary of spray coverage to determine the width of coverage ^34-36^. The coverage rates are listed in Table 5.

**Table 5.**
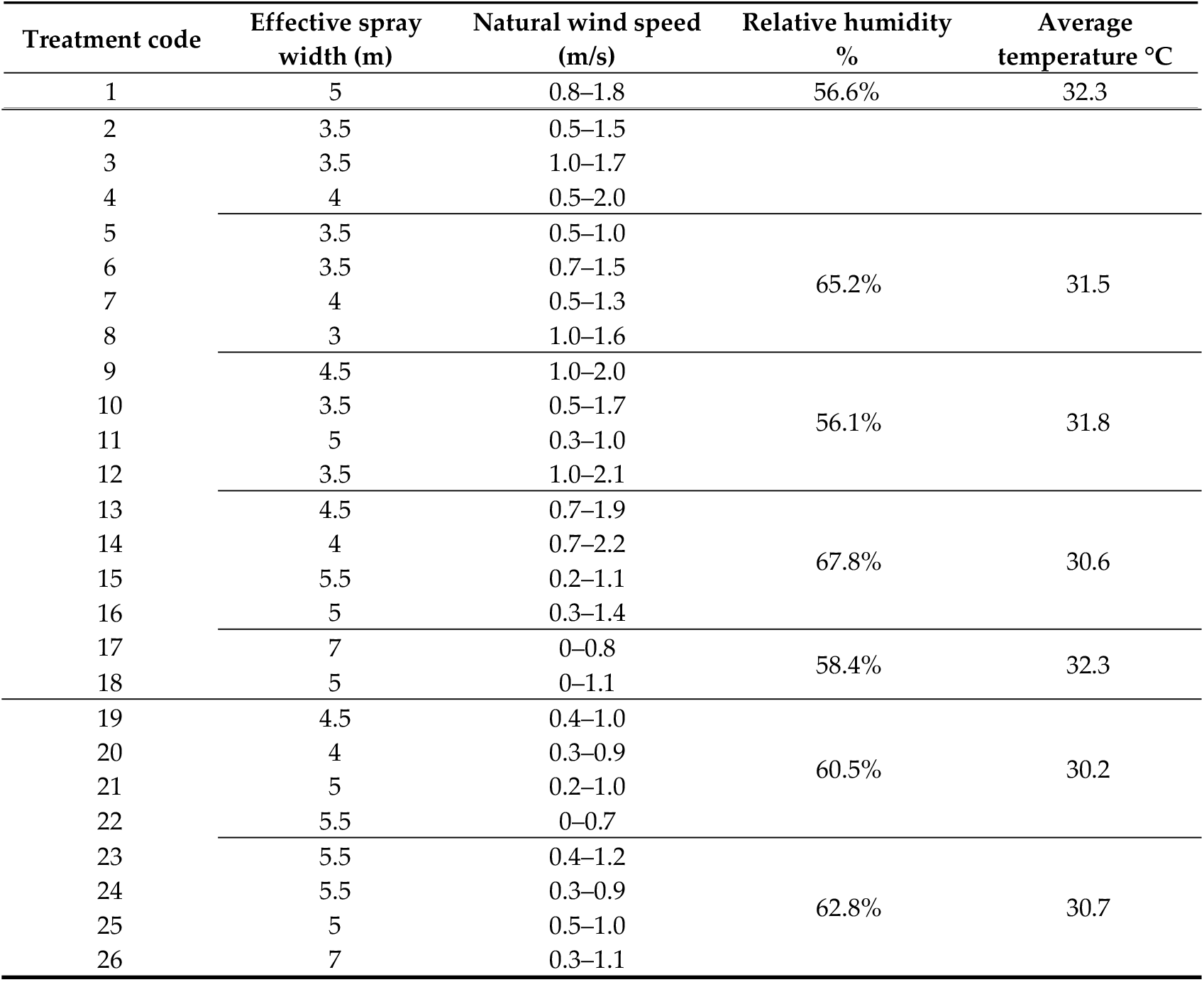
Effective spray width for different treatments.

The field efficacy was calculated according to the following formula^37^:

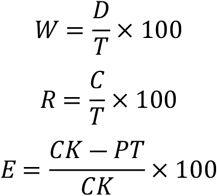

where W represents the blight heart rate, %; D denotes the number of blight heart plants investigated; T indicates the total number of plants investigated; R represents the rolling leaf rate, %; C denotes the number of rolling leaves investigated; T indicates the total number of leaves investigated; E represents the control efficacy, %; CK denotes the blight heart rate or rolling leaf rate in the blank control area after spraying; and PT indicates the blight heart rate or rolling leaf rate in the spraying area.

### 2.4. Basic Information Record

The crop growth and weather conditions, which are listed in Table 6, were recorded.

**Table 6.**
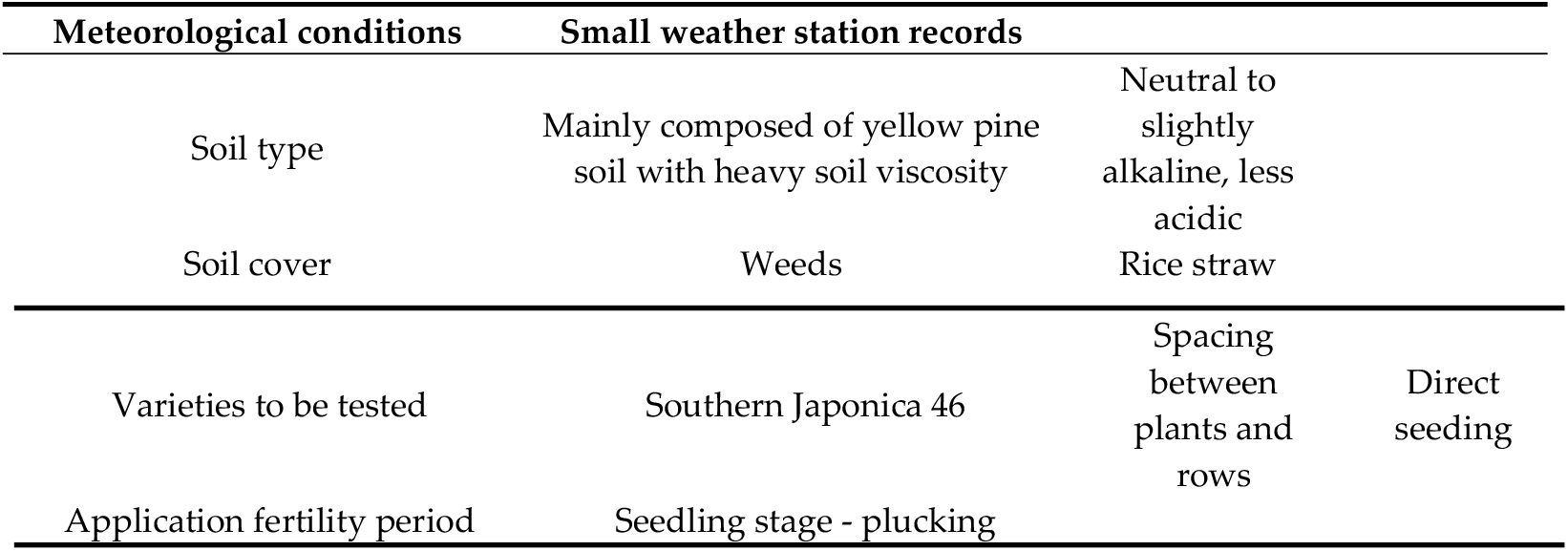
Basic information record form.

| Meteorological conditions | Small weather station records |  |
| --- | --- | --- |
| Soil type | Mainly composed of yellow pine soil with heavy soil viscosity | Neutral to slightly alkaline, less acidic |
| Soil cover | Weeds | Rice straw |

**Table 6.** Basic information record form.
|  |  |  |  |
| --- | --- | --- | --- |
| Varieties to be tested | Southern Japonica 46 | Spacing<br>between<br>plants and<br>rows | Direct<br>seeding |
| Application fertility period | Seedling stage - plucking |  |  |

## 3. Results and Analysis

### 3.1. Effects of different adjuvants on the spread of rice

The spreading characteristics shown in Figure 5 clearly followed a hierarchy: Activator 90 > TM07 > Sylgard 309 > Aeromate 480 > Silwet L-77 > LI-700. This pattern is strongly correlated with the surface tension data in Table 1. Activator 90 and Silwet L-77, with the lowest surface tensions (20.5 mN/m and 18.9 mN/m, respectively), exhibited immediate spreading within 1–2 seconds, which is consistent with their surfactants rapidly reducing the liquid surface tension. In contrast, LI-700, with the highest surface tension (30.5 mN/m) and viscosity (2.80 mPa·s), had the poorest spreading performance, requiring 4–5 minutes to achieve limited coverage. The delayed spreading of TM07 and Sylgard 309, despite their moderate surface tensions (23.8 and 22.0 mN/m), suggests that their spreading mechanisms involve more complex interfacial interactions rather than immediate surface tension reduction.

**Figure 5.**
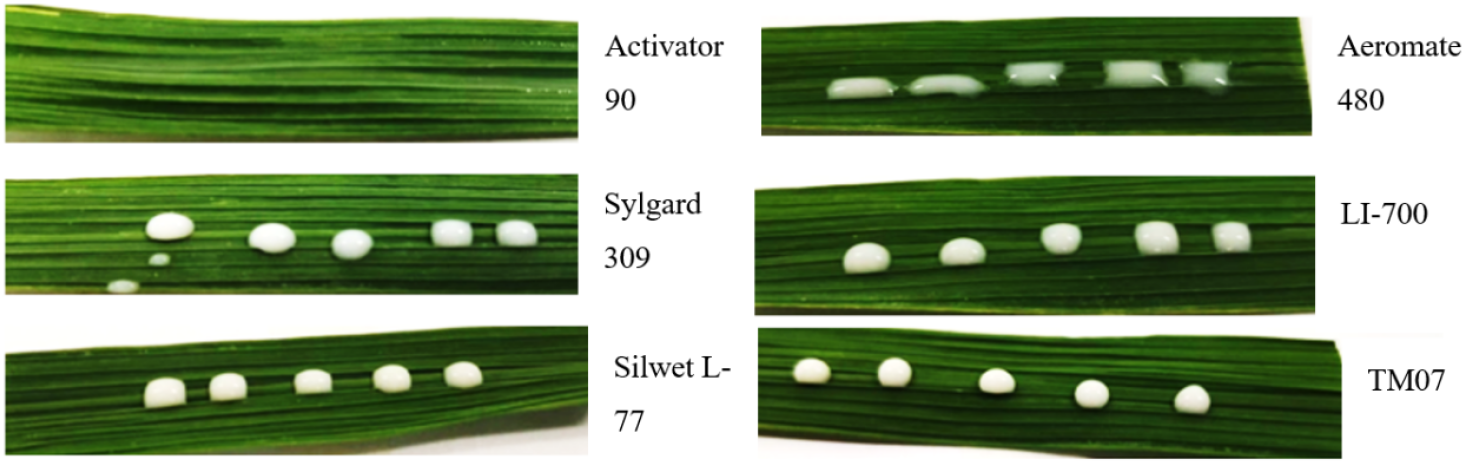
Spreading effect of different adjuvants on rice leaves. **Note:** Each image was taken when the droplet reached a visually stable spreading state for the corresponding adjuvant.

### 3.2. Droplet deposition of different adjuvants

The deposition results in Table 7 reveal that adjuvants with balanced physical properties achieved optimal performance. Activator 90, with low surface tension (20.5 mN/m) and moderate viscosity (1.05 mPa·s), achieved the highest deposition density (30.0 drops/cm^2^) and lowest coefficient of variation (44.6%), indicating that its properties facilitated both adequate droplet formation and stable flight characteristics. Aeromate 480, despite having the second-highest viscosity (2.50 mPa·s), achieved high deposition density (42.3 drops/cm^2^) but with high variability (CV=60.1%), suggesting that its drift-reducing properties came at the cost of distribution uniformity. LI-700, which combined high viscosity and high surface tension, had the poorest performance (CV=99.1%), because these properties likely hindered both droplet fragmentation and target surface adhesion.

**Table 7.** Droplet deposition density and coefficient of variation in the upper crop layer for different adjuvant treatments.

| Treatment code | Machine type | Operating height | Operating speed | Application rate | Pesticides and adjuvants | Deposition density (drops/cm<sup>2</sup>) | Coefficient of variation (%) |
| --- | --- | --- | --- | --- | --- | --- | --- |
| 1 | T30 Plant<br>Protection<br>UAV | 2.0 m | 4 m/s | 15 L/ha | Pesticides<br>+Silwet L-77 | 19.7±6.49 | 56.2±19.86 |
| 2 |  |  |  |  | Pesticides<br>+Sylgard<br>309 | 27.0±8.23 | 54.9±19.39 |
| 3 |  |  |  |  | Pesticides<br>+Activator<br>90 | 30.0±9.14 | 44.6±15.78 |
| 4 |  |  |  |  | Pesticides<br>+TM07 | 42.1±14.26 | 48.3±17.04 |
| 5 |  |  |  |  | Pesticides<br>+LI-700 | 28.6±9.60 | 99.1±35.07 |
| 6 |  |  |  |  | Clearwater | 39.5±12.49 | 109.0±38.44 |
| 7 |  |  |  |  | Pesticides | 44.0±13.75 | 95.5±33.67 |
| 8 |  |  |  |  | Pesticides<br>+Aeromate<br>480 | 42.3±13.40 | 60.1±21.20 |

The sediment distribution of the droplets with and without adjuvants at different flight speeds at a spraying rate of 15 L/ha and a flying height of 2.5 m is shown in Figure 6. At the same flying height and spraying rate, the addition of adjuvants significantly increases the droplet sediment density, indicating that the addition of adjuvants facilitates droplet sediment formation. In addition, at different flight speeds, the addition of adjuvants can also increase the droplet sedimentation density, indicating that adjuvants positively affect pesticide spraying with different operating parameters.

**Figure 6.**
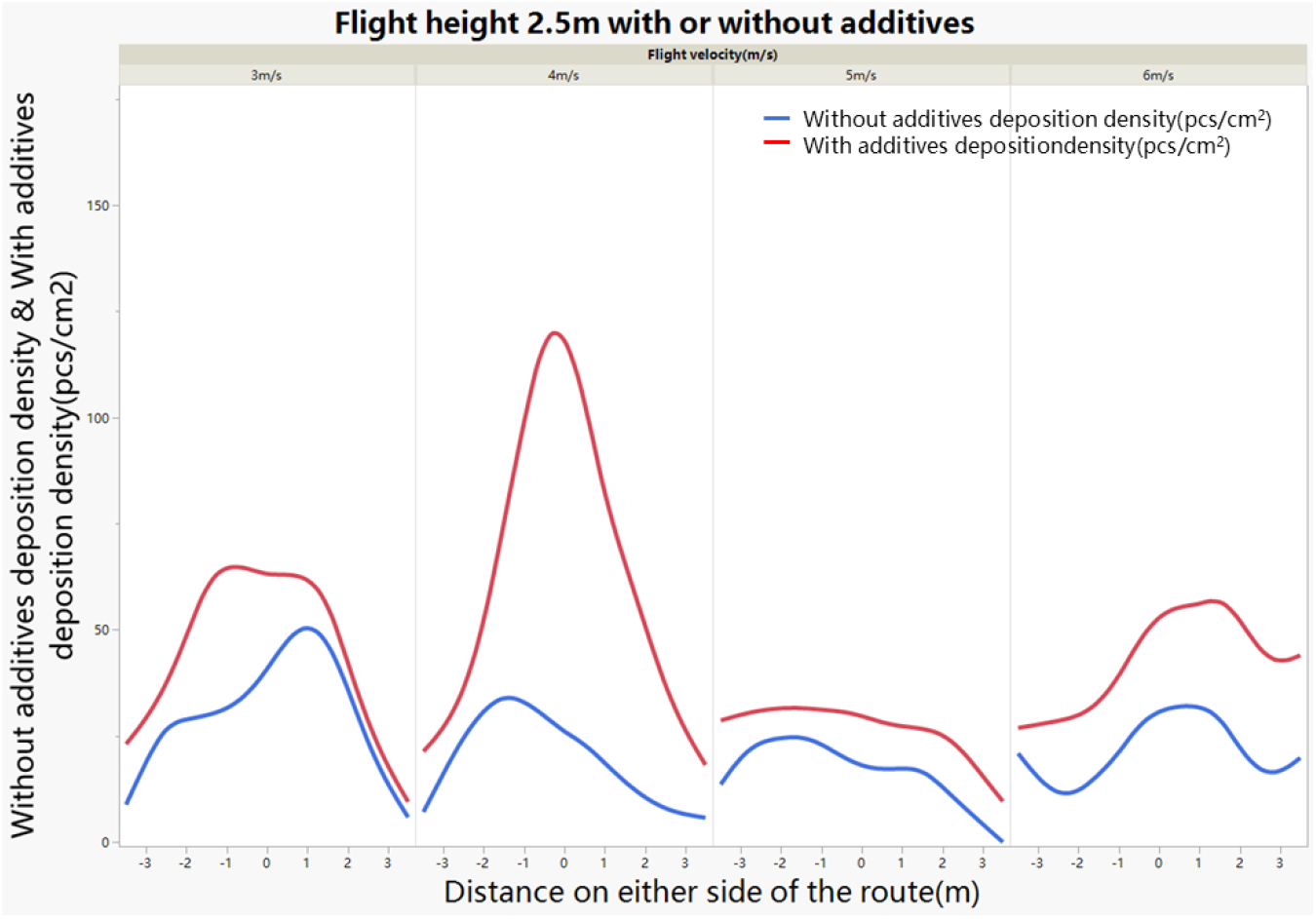
Comparison of deposition distribution of adjuvant-free and adjuvant-treated aerosol droplets.

These results reveal that the addition of spray adjuvants can significantly affect the droplet deposition pattern on rice crops. Activator 90 resulted in the lowest coefficient of variation (44.6%), indicating more uniform coverage, which is critical for consistent pest control. This finding is consistent with the findings of Acevedo et al.,^39^ who reported that nonionic surfactants can reduce surface tension and facilitate uniform dispersion of fine droplets during aerial application. Although Aeromate 480 had a high droplet density (42.3 drops/cm^2^), its relatively high variation (60.1%) suggests that the droplets may have clustered or drifted because of less efficient spreading. Interestingly, even without any adjuvant, clearwater treatment resulted in a high deposition density (39.5 drops/cm^2^), but the associated high variation (109.0%) highlights the importance of uniformity, not just quantity, in effective pesticide application. Overall, while both droplet quantity and consistency are crucial, adjuvants such as Activator 90 and TM07 appear to strike a better balance between the two, increasing both deposition density and distribution uniformity under UAV spraying conditions.

When plant protection UAVs are used for pest and disease control, droplet deposition coverage, droplet density, and sedimentation uniformity significantly affect the effectiveness of pest control^38^. Previous studies have shown that spraying uniformity is challenging with UAVs for crop protection, because factors such as rotor wind fields and nozzle positions can affect the uniformity of spraying^40^. The results of this study revealed that the use of adjuvants further improved the uniformity, which significantly affected the effectiveness of pest and disease control. Therefore, when UAVs are selected for spraying operations, suitable spray adjuvants should be chosen as much as possible.

### 3.3. Spraying effect of different adjuvants

A variance analysis was performed on the various factors in the experiment, and a Duncan multiple range test was used for significance testing. The results are shown in Table 8. The effects of the adjuvant and the application rate per ha on the spray width are extremely significant, whereas the effect of height on the spray width is significant, and the effect of speed on the spray width is not significant. Under the same spraying conditions, using an adjuvant and increasing the application rate per ha can significantly improve the spraying effect. Within a specific range, the spraying speed has a relatively small effect on the spraying effect. Owing to the limited application rate and adjuvant treatment, the degrees of freedom for these factors were minimal. Although significant differences were observed, further studies with more treatment levels are needed to confirm these effects.

**Table 8.** Analysis of variance table for effective spray widths.

| Source of variance | Freedom | Mean square sum | F value | Significance P value | Significant level |
| --- | --- | --- | --- | --- | --- |
| Height A | 3 | 1.7 | 3.8 | 0.03 | * |
| Speed B | 3 | 1.0 | 2.3 | 0.12 | Insignificant |
| Application rate C per ha | 1 | 10.6 | 23.8 | <0.001 | ** |
| Adjuvants | 1 | 8.3 | 18.6 | <0.001 | ** |
| Inaccuracy | 17 | 0.45 |  |  |  |

According to Figure 7, the spray width increases with increasing application rate, likely because of the greater volume of liquid being discharged, which allows the droplets to cover a broader area. Furthermore, the addition of spray adjuvants appears to further improve this effect, potentially by improving droplet adhesion and retention on crop surfaces. This approach may facilitate more effective deposition and spread of the spray solution, thereby improving coverage and pest control performance. The experimental results also revealed that the spraying speed has a minimal effect on the spraying range; that is, at a specific spraying amount and height, the speed of spraying does not play a decisive role in the change in the spraying range. The results of the experiments also revealed that as the spraying height increased, the spraying height first increased but then decreased, and the maximum spraying range occurred at a height of 2.5 m. As the spraying height increased, the vertical velocity of the spray particles increased, thereby increasing the spraying range. However, as the spraying height continues to increase, the frictional resistance between the spray particles and the air increases, causing the spray to drift upwards, resulting in a gradual decrease in the spraying range. Under the conditions tested in this study, the maximum spraying width occurred at a spraying height of 2.5 m, an application rate of 22.5 L/ha, and with the addition of an adjuvant.

**Figure 7.**
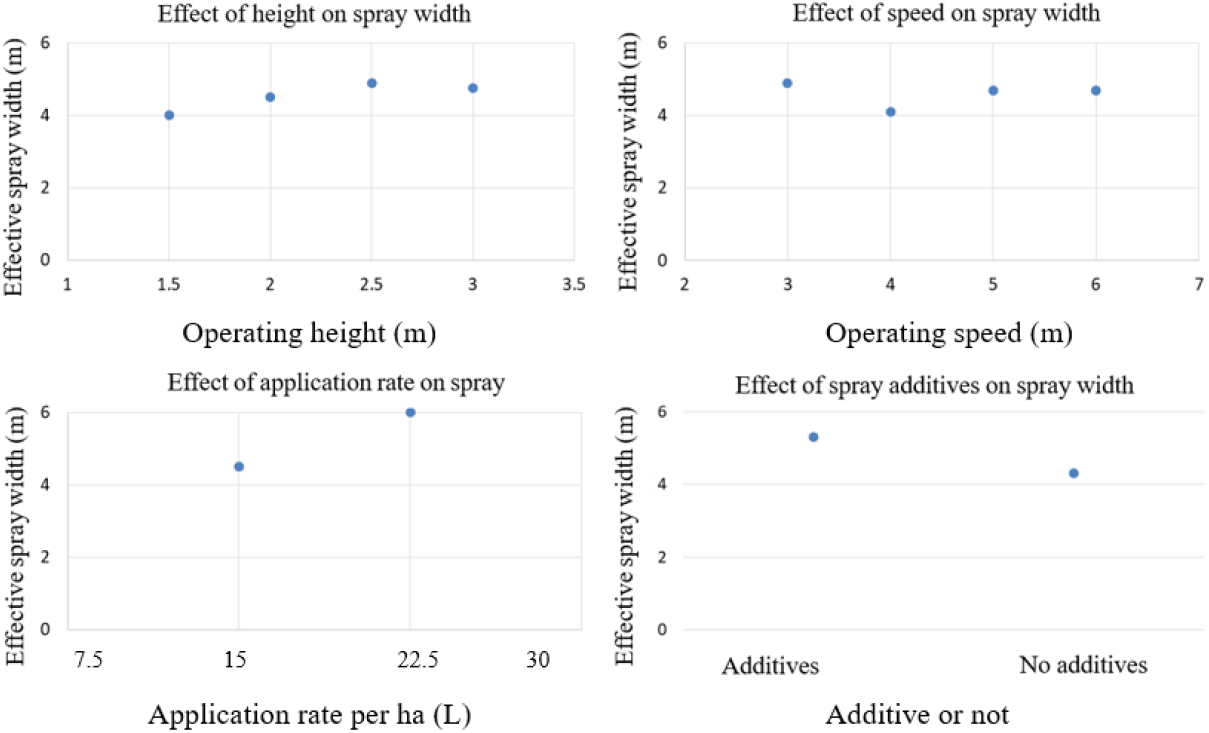
Effects of various factors on spray width.

### 3.4. Field Control Efficacy of Different Adjuvants against Rice Stem Borers

The field efficacy results in Table 9 reveal how physical properties ultimately translate to biological effectiveness. The excellent performance (87% control efficacy) of Activator 90 stems from its optimal combination of low surface tension for improved leaf coverage and adequate viscosity for deposition retention. The moderate performance of Sylgard 309 (78% efficacy), despite its favourable surface tension (22.0 mN/m), may be attributed to its oil-based composition potentially affecting active ingredient availability. The lower efficacies of LI-700 and Aeromate 480 (77% each) reflect the limitations of their extreme physical properties—while high viscosity reduces drift, it may also impede droplet penetration and redistribution on target surfaces.

**Table 9.**
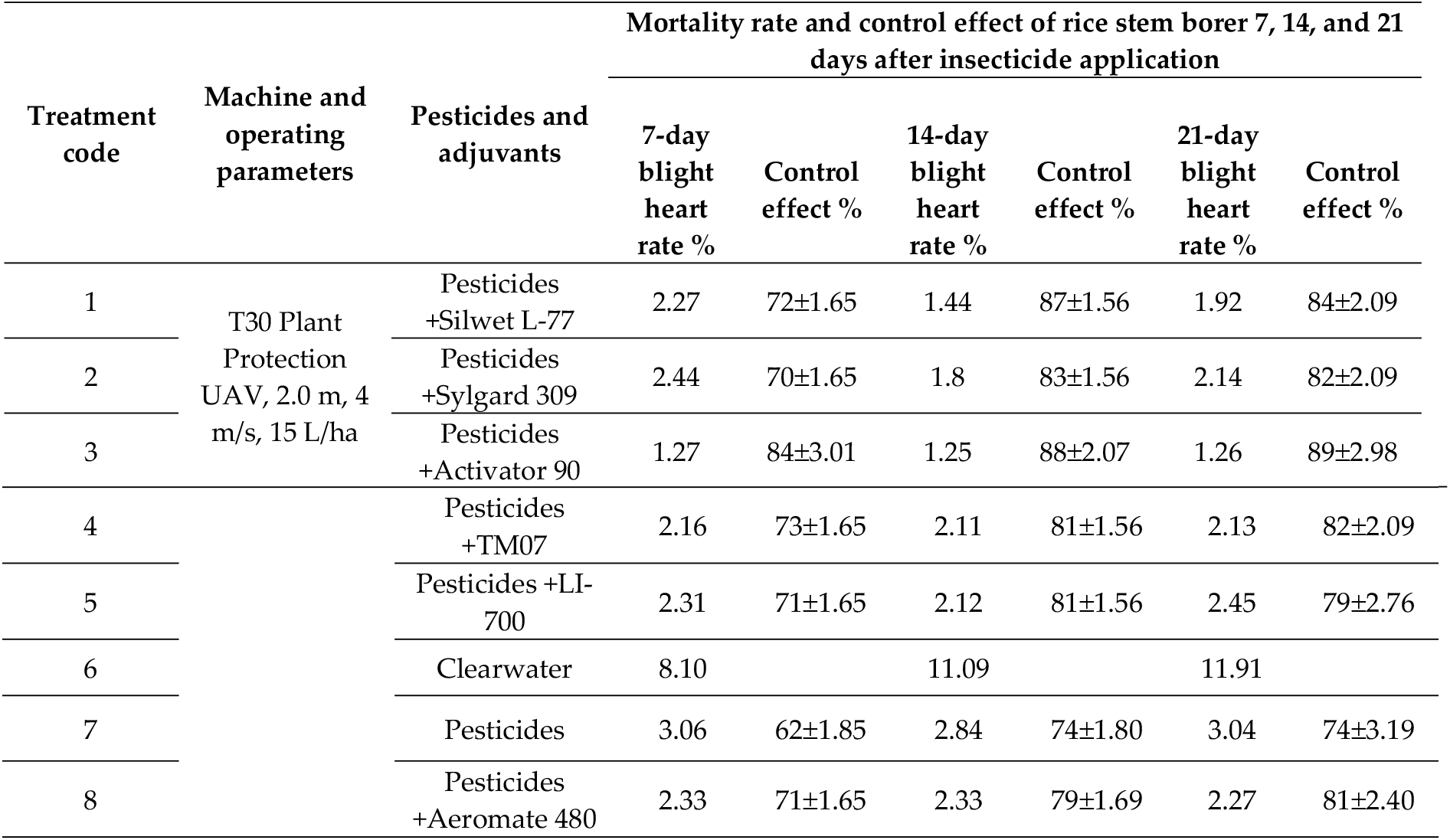
Effectiveness of prevention and treatment.

The results clearly indicate that the addition of spray adjuvants increases the control efficacy of chlorantraniliprole against rice striped stem borers. Activator 90 stood out, with an average control effect of 87%, which was significantly greater than that of the other treatments, which is consistent with its excellent droplet spreading and uniform deposition observed in earlier sections. These findings support previous findings by Silva et al.,^41^ who reported that improved spreading and retention of active ingredients on plant surfaces can increase insecticidal activity. In contrast, the pesticide-only treatment (Treatment 7) and clearwater treatment (Treatment 6) had the lowest control effects, indicating that without adjuvants, pesticide performance is hindered, likely because of reduced adhesion and penetration. Although Silwet L-77 and TM07 also had relatively strong control effects (81% and 79%, respectively), their delayed spreading behaviour may limit rapid absorption of the pesticide, especially under conditions of high humidity or rain. Furthermore, while Aeromate 480 and LI-700 achieved moderate efficacy (approximately 77–79%), the higher variability in droplet deposition observed with LI-700 may have contributed to inconsistent coverage and reduced effectiveness. Overall, these results highlight the importance of choosing appropriate adjuvants on the basis of not only their deposition characteristics but also their ability to increase pesticide uptake and pest mortality.

An issue worthy of further exploration is that deposition density does not exhibit a straightforward positive correlation with final control efficacy. For example, the treatments with the highest deposition densities in this study (Aeromate 480 at 42.3 drops/cm^2^; pure pesticide at 44.0 drops/cm^2^) did not yield optimal control outcomes. Conversely, Activator 90—which had moderate deposition density (30.0 drops/cm^2^) but the highest spray uniformity (CV 44.6%)—had the highest control efficacy (87%). CV 44.6%) had the highest efficacy (87%).

This outcome highlights the decisive role of droplet distribution uniformity in pest control. Excessively high coefficients of variation (such as 95.5% for the pesticide-only treatment) indicate clustered droplet distribution within the canopy, creating extensive uncovered or inadequately covered ‘safe zones’ where pests can survive and reproduce. Conversely, Activator 90 significantly improves deposition uniformity, ensuring continuous and complete effective coverage of the target area. This maximizes the elimination of pest refuges, achieving more thorough population control.

## 4. Conclusion

Indoor experiments in this study revealed that when different adjuvants were added to the pesticide solution and droplets were applied to the rice leaves, significant differences in the spreading characteristics and spreading time were observed. On the basis of their spreading performance, the overall ranking was as follows: Activator 90 > TM07 > Sylgard 309 > Aeromate 480 > Silwet L-77 > LI-700. This ranking is closely related to the physical properties of the adjuvants, particularly surface tension, which is consistent with recent research indicating that low surface tension adjuvants (e.g., Activator 90) can rapidly disrupt the interfacial energy between the droplet and the leaf surface, facilitating fast spreading.

Field spray width tests revealed that the spray width increased with increasing application volume, and the addition of adjuvants effectively increased the spray width, confirming that adjuvants facilitate pesticide deposition by regulating the physicochemical properties of the spray liquid (e.g., reducing surface tension and moderately increasing viscosity). The spray width first increased but then decreased with increasing flight height, reaching its maximum at 2.5 metres. This phenomenon is related to the interaction between the UAV downwash airflow and the droplet sedimentation trajectory. Recent studies have also indicated the existence of an optimal height range to balance penetration and drift risk. The optimal parameter combination for achieving the maximum spray width was a flight height of 2.5 m, an application volume of 22.5 L/ha, and the addition of adjuvants.

Analysis of the droplet deposition density and the coefficient of variation (CV) of the upper crop layer elucidated the distribution patterns of deposition with and without different adjuvants. Overall, the data revealed that adjuvants significantly reduced the CV and improved the uniformity of the droplet distribution. The Activator 90 treatment resulted in the smallest CV and the best deposition effect, followed by those of Sylgard 309 and TM07. There was no significant difference in the improvement effect between Silwet L-77 and Aeromate 480, whereas LI-700 performed the worst. These findings highlight that in modern precision pesticide applications, deposition uniformity is as crucial as, if not more critical than, absolute deposition density because uniform coverage decreases the number of control blind spots. This finding aligns closely with recent conclusions on “effective coverage”.

Field efficacy trials confirmed that the addition of adjuvants significantly increased the control efficacy of chlorantraniliprole against the rice stem borer. During the 21-day investigation period, the average control efficacy for adjuvant-treated plots was 77%–87%, whereas it was only 70% for the treatment without adjuvant. The synergistic effects of the different adjuvants were ranked as follows: Activator 90 > Silwet L-77 > TM07 > Sylgard 309 > Aeromate 480 = LI-700. This ranking is largely consistent with the ability of adjuvants to improve deposition uniformity, further confirming that ensuring uniform deposition by optimizing physical properties is the main pathway to achieving stable and highly effective field performance.

## Data Availability Statement

The datasets used and/or analysed during the current study are available from the corresponding author upon reasonable request.

## Statement

Permission was obtained from the relevant authorities and stakeholders to conduct this study in the agricultural field.

